# Influence of temperature on growth and development of dictyostelid slime moulds and its implication on the evolution of cold-tolerance

**DOI:** 10.1101/2022.09.19.508496

**Authors:** Hidenori Hashimura, Kei Inouye

**Affiliations:** Department of Botany, Graduate School of Science, Kyoto University, Kyoto 606-8502, Japan; Department of Biological Sciences, Graduate School of Science, Osaka University, 1-3 Yamadaoka, Suita, Osaka, 565-0871, Japan; RIKEN Center for Biosystems Dynamics Research (BDR), 6-2-3 Furuedai, Suita, Osaka, 565-0874, Japan; Graduate School of Arts and Sciences, University of Tokyo, 3-8-1 Komaba, Meguro, Tokyo 153-8902, Japan

## Abstract

Environmental temperature is a major determinant of microbial life. Dictyostelids are soil amoebae capable of multicellular social behaviour upon starvation. They inhabit in a variety of environments from the tundra to the tropics, but how they have adapted to environmental temperature remains largely unknown. In this study, the effects of temperature on the growth and multicellular development of 36 dictyostelid species (58 strains/isolates) were examined. More than half of the species showed maximal growth and normal development at 28°C or above, whereas some could grow and develop at 4°C, or even at 0°C. Many of the isolates examined were from areas with temperatures far lower than their preferred range over a large part of the year. There was a significant correlation between thermal characteristics and phylogeny. Over 150 known dictyostelid species are divided into several taxonomic groups. Our phylogenetic analysis indicated that cold-tolerance evolved independently in major clades, most prominently in group 4 (genus *Dictyostelium* according to the new classification by Sheikh *et al*.), which contains many species that are often found in subarctic regions. These results suggest that ancestors that have acquired cold-tolerance expanded their ranges into cooler areas where they could proliferate and develop during summer and survive the severe winter.

## Introduction

Temperature is an important factor affecting biological activity, and it is a major determinant of the global distribution of organisms. In general, organisms adapt to environmental temperature by genetic alteration or acclimation [1]. In recent years, it has become widely appreciated that closely related species share similar thermal traits and geographic distribution patterns [2]. Since the influence of phylogeny on thermal traits and species distribution provides essential information to understand the evolution of thermal adaptation and how the current distribution of species were formed, phylogenetic comparative studies have been conducted in various organisms such as ectothermic animals and invertebrates [3],[4]. However, studies focusing on such relationships have been scarce in eukaryotic microbes.

Dictyostelids (cellular slime moulds) are one of the most studied soil-living amoebae mainly because of its experimental tractability. The amoebae usually feed on bacteria and proliferate, but they show the unique development when they deplete the food source. Starved amoebae aggregate to form multicellular fruiting bodies consisting of spores and a stalk which supports the spore mass. While stalk cells all die, spores are resistant to environmental stress (*e.g.*, temperature and desiccation) and survive long periods of starvation by remaining dormant until improvement of circumstance or being transported to new feeding sites by various animals such as small invertebrate or other agents [5]. Over 150 species have been described so far. At present, they are classified into four major clades and several minor clades based on SSU rDNA and protein sequence data [6]–[9].

While climate conditions such as temperature and water availability generally affect species richness [10], dictyostelids have been isolated from a wide variety of climate zones including the tundra and the tropics [11]. Several literatures have reported thermal traits that are seemingly adaptive to their habitats. Species isolated from the tropics and subtropics can grow and develop optimally at >28°C [12]–[15], which is above the optimal range (22–25°C) for activity of the other species including *Dictyostelium discoideum* [14]. On the other hand, isolates from the cold region prefer the lower temperature, and their growth and development are inhibited at high temperature [12],[16],[17]. These examples may suggest that dictyostelids expand their habitats across a wide range of climate zones through evolution. However, our understanding of the thermal characteristics in dictyostelids is still far from complete, because it is only based on a small number of species [13],[18]. This lack of systematic studies makes it difficult to investigate their relationship with the phylogeny and to understand their possible effects on the global distribution of the dictyostelids.

In this study, we examined how temperature affects the growth and development of various dictyostelid species covering their entire taxonomic breadth. We confirmed the diversity of thermal characteristics between species such as the preference of high temperature by many species, and found unexpected ability to grow and develop at 0°C in some species. Comparative analysis of the temperature effects revealed the influence of phylogeny on thermal tolerance and preference. We will discuss possible scenarios for the divergence of thermal characteristics in the evolutional history of dictyostelids, and how its diversity may be related to the formation of their current geographical distribution.

## Materials and methods

### Species identification

The dictyostelids used in this study were isolated from soil samples collected at various sites in Japan, or obtained from ATCC, Dicty Stock Center and NBRP-nenkin, or laboratory stocks. These are listed in S1 Table, in which species names according to the new nomenclature by Sheikh *et al.* (2018) [19] are given along with the old names. In the other part of this paper, basically old names are used (new names in parentheses). In the figures and tables, a 4-letter abbreviation of the old names (see S1 Table) is also used. Detailed information of every strain is compiled in S2 Table. Species identification of the local isolates was based on morphology and SSU rDNA sequences. The species names that gave the highest match scores in blastn searches against the NCBI nucleotide database were adopted. For the species obtained from the stock centers and laboratory stocks, the designated species and strain names are used in the present paper. There are two instances of disagreement between the designated species name and that by the rDNA sequence; *D. mucoroides* no.7 [20] is *D. clavatum* according to SSU rDNA, and *P. palladium* (*Heterostelium pallidum*) PN500 [21] is *P. album* (*Heterostelium album*) [22]. It should be noted that the isolates identified as *D. brefeldianum* in this study is most likely identical to *D. mucoroides* in the papers cited in this paper [23]. For sequencing, the SSU rDNA or SSU + ITS region of rDNA was amplified by PCR using primers described previously [24] or SSU start F2 (ACTGGTTGATCCTGCCAGTAG) and ITS R (CTTACTGATATGCTTAAGTTCAGCGG). The amplified DNA was purified using QIAquick gel extraction kit (QIAGEN) or Wizard PCR purification kit (Promega), and then directly sequenced using the same and internal primers on an ABI 3130xl sequencer.

### Culture

All species were cultured in association with the gram-negative bacteria, *Klebisiella aerogenes*. The bacteria was pregrown in shaken suspension of 100 ml liquid SM medium (KH_2_PO_4_ 4.4 g, Na_2_HPO_4_·12H_2_O 2.0 g, D-Glucose 7.5 g, Bacteriological Petone (Difco) 10 g, Yeast extract (Difco) 1.0 g, MgSO_4_·7H_2_O 1.0 g, H_2_O 1,000 ml [25]) for 24 hours at 22°C. The bacteria were harvested by centrifugation at 5,000 rpm for 5 minutes at 4°C, resuspended in 10 ml of 20 mM KK2 phosphate buffer, pH 6.0, and stored at 4°C until use. Dictyostelid cells used in temperature experiments were pregrown on the LP agar plate (0.1% lactose, 0.1% Bacteriological Petone, 20 mM KK2 phosphate buffer, pH 6.0) with the bacteria at 22°C, or at 16°C for *D. septentrionalis* (*D. septentrionale*) which does not grow at 22°C [17], for 24 to 48 hours before the start of experiments at indicated temperatures. For shaking culture of the dictyostelids, spores or amoebae were inoculated in 9.3 ml of KK2 buffer to which 0.7 ml bacteria stock solution had been added, and incubated at 22°C in shaken suspension at 175 rpm. Some species that did not grow well in suspension (S3 Table) were grown on SM agar plates as follows: 5×10^5^ cells were suspended in 100 μL KK2 buffer to which 50 μL bacteria solution had been added. The suspension was spread on SM agar plate and incubated at 22°C for 24 to 48 hours. Cells were harvested before clearing of the bacterial lawn and washed three times by centrifugation at 1,500 rpm for 1 minute, and resuspended in KK2 buffer.

### Growth experiments

Temperature effects on growth were monitored using two conditions; circular spreading on a bacterial lawn on nutrient-rich SM agar, and linear spreading on a bacterial streak on nutrient-poor LP plate. For the former, 50 μL of the bacterial suspension was diluted in 100 μL of KK2 buffer and spread on SM agar plates, then incubated at 22°C for 24 hours. A 10 μL droplet containing ca. 10^4^ of Dictyostelid cells was deposited on the centre of the pre-incubated, fully grown bacteria. For the latter method, diluted bacterial suspension was streaked (3 or 7 cm long depending on species) on LP agar plate, and incubated at 22°C overnight before inoculation of cells at one end of the streak. Plates were incubated over a period of a month or longer at 0, 4, 10, 16, 22, 28, 31, 34, or 37°C. For incubation at 0°C, plates were put in a plastic box or bag and buried in crushed ice (ca. 20 kg) in a large styrofoam box, which was kept in a cold room kept at 4°C. The temperature around the plates was occasionally checked. Plaque size on the bacterial lawn or the advancement of the feeding edge along the streaks was measured daily (weekly for 0°C) (S1 Fig). The growth ability at each temperature was evaluated from these values (cm/day). The temperature giving the highest plaque growth rate was taken as the optimal growth temperature. For evaluation of temperature dependence, relative growth rate, normalised to the growth rate at the optimal growth temperature, was used. Since the nutrient-rich and nutrient-poor conditions gave very similar relative growth rate values at each temperature for all strains, the data of the two conditions were combined and the means were used in the analysis. Throughout the study, we used relative growth rate normalised to the maximum growth rate for each isolate. The raw growth rates as determined in two-membered culture substantially differ among species, being very high in species of the violaceum and group 4 clades and low in those of groups 2A and 3. However, we consider that the difference will not be large, or may even disappear, in natural habitats, because species that grow slowly in common laboratory conditions are not necessarily rare in nature. We have often isolated such slow growers as *Acytostelium subglobosum* and *D. minutum* (*Raperostelium minutum*) along with fast growers such as *P. violaceum* and *D. purpureum* from a few grammes of soil samples collected at the same site in several successive years, without any sign of the slow growers’ diminishing.

### Temperature effects on development

To assess the temperature dependence of development, a 5 μL droplet containing 10^5^ or 5×10^5^ cells was deposited on 2% [w/v] autoclaved water or KK2 agar plates (Difco Bacto agar in distilled water or KK2 buffer). For *D. polycarpum* (*Synstelium polycarpum*), water agar containing 0.5% [w/v] charcoal was used to encourage fruiting. Plates were incubated over a period of a month or longer at 0, 4, 10, 16, 22, 28, 31, 34, or 37°C. Development of the cells was monitored daily (weekly for 0°C) and the developmental ability at each temperature was assessed from the morphology of aggregation and multicellular structures formed. In *D. polycephalum* (*Coremiosteium polycephalum*), slugs kept migrating for weeks and seldom made fruiting bodies at any temperature. The development of this species was therefore considered complete upon the formation of migrating slugs.

### Data analysis

For comparative analysis, several measures of thermal characteristics were defined and calculated from the growth rate vs. temperature (G-T) relationship for each strain/isolate. Optimal growth temperature is the temperature giving the highest rate of growth. Temperatures giving x% maximal growth were directly calculated from G-T relationship. On the other hand, optimal growth temperature and lower and upper thermal tolerance limits were estimated from the G-T relationship (abbreviated as LLx and ULx, respectively) using its 1^st^, 2^nd^, and 3^rd^ order moments (equations in S1 Appendix), because it was impracticable to set the number of temperature points sufficiently large for direct determination of these variables due to the large number of strains/isolates to be examined. Mean growth temperature was defined as the 1^st^ order moment of the G-T relationship.

### Phylogenetic analysis

The published phylogeny based on SSU rDNA sequence [6],[7] was used for analysis of the 36 representative species which are included in this phylogeny. For analysis with additional strains (S2 Fig), local trees based on SSU rDNA sequences were constructed and embedded in the main tree after standardisation of the branch lengths. To estimate phylogenetic signal, we calculated Pagel’s λ [26], Blomberg’s K [27] and Moran’s I [28], using the R packages phytools [29], phylobase [30], and adephylo [31]. Ancestral state reconstruction was performed with the contMap function in the phytools package.

### Acquisition and handling of climate data

Climate data of the locations of sampling sites were obtained from WorldClim (http://www.worldclim.org) [32], which provides various climate variables and altitude data at 30 arc-second resolution (ca. 930 m at the equator). Many of the strains obtained from culture collections lack precise location of isolation site. Efforts were made to narrow the area of possible locations mainly on the basis of the place names and altitudes given in the original report. If altitude is missing, the possible area was limited to plus-minus 500 m in altitude from the centre of the named location. Two strains of which the isolation site could not be identified were excluded from the analysis concerning climate. For the locations encompassing more than one 30-arcsec cells, minimum, maximum, and mean of each monthly average temperature were obtained, and shown in the graphs of Fig 5 as a range of temperature. The monthly average temperatures and the global coordinate values of the locations of isolation site are summarised in S4 Table. Data acquisition from the WorldClim site and data extraction were conducted using R 3.2.1, and data processing with perl scripts.

## Results

### Thermophicility and cold-tolerance of dictyostelids

We examined the effects of temperature (0 – 37°C, 9 temperature points) on growth and development of 36 species (58 strains/isolates) covering the taxonomic range of dictyostelids (S1 Table). The growth of each species was quantified by daily measurements of the spread on bacterial lawns or streaks on nutrient agar plates. Fig 1 shows examples of growth curves at various temperatures (S1 Fig for growth curves of other species/strains under nutrient-rich and nutrient-poor conditions). The rate of growth was naturally dependent on temperature, and the maximum growth in 24 hours at each temperature was regarded as the growth rate at that temperature. The temperature giving the highest growth rate will be called the optimal growth temperature of that species. The development of each species at various temperatures was monitored by incubating pregrown cells on non-nutrient agar plates. The development of cells inoculated at one end of bacteria streaks on nutrient-poor LP plates was also examined. These two conditions gave similar results unless otherwise noted. The ability of development at each temperature was categorised into (i) formation of fruiting bodies, (ii) formation of aberrant multicellular structures (examples shown in Fig 2) (iii) arrest after aggregation, and (iv) absence of aggregation.

**Fig 1.**
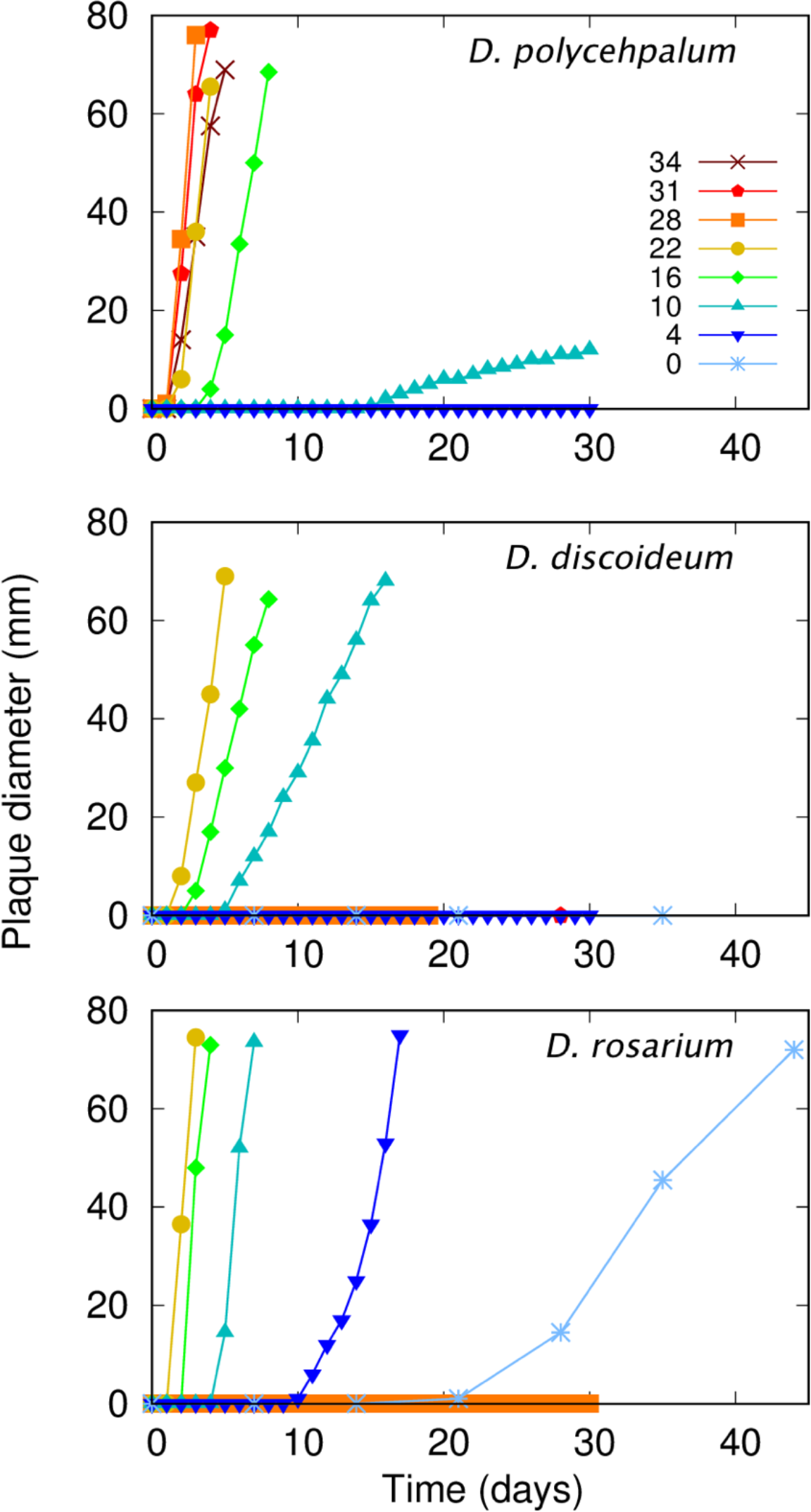
Growth curves of three species at various temperatures. Growth curves of *D. polycephalum* (*C. polycephalum*) 1JND1, *D. discoideum* NC4, *D. rosarium* W S689 are shown. A 10 microlitre droplet of cell suspension was deposited on a bacterial lawn on nutrient-rich agar plates, and incubated at 0 – 37°C. The diameter of the plaque was measured daily (4 – 37°C) or weekly (0°C). No growth was observed in any species at 37°C, which is not shown on the graphs.

**Fig 2.**
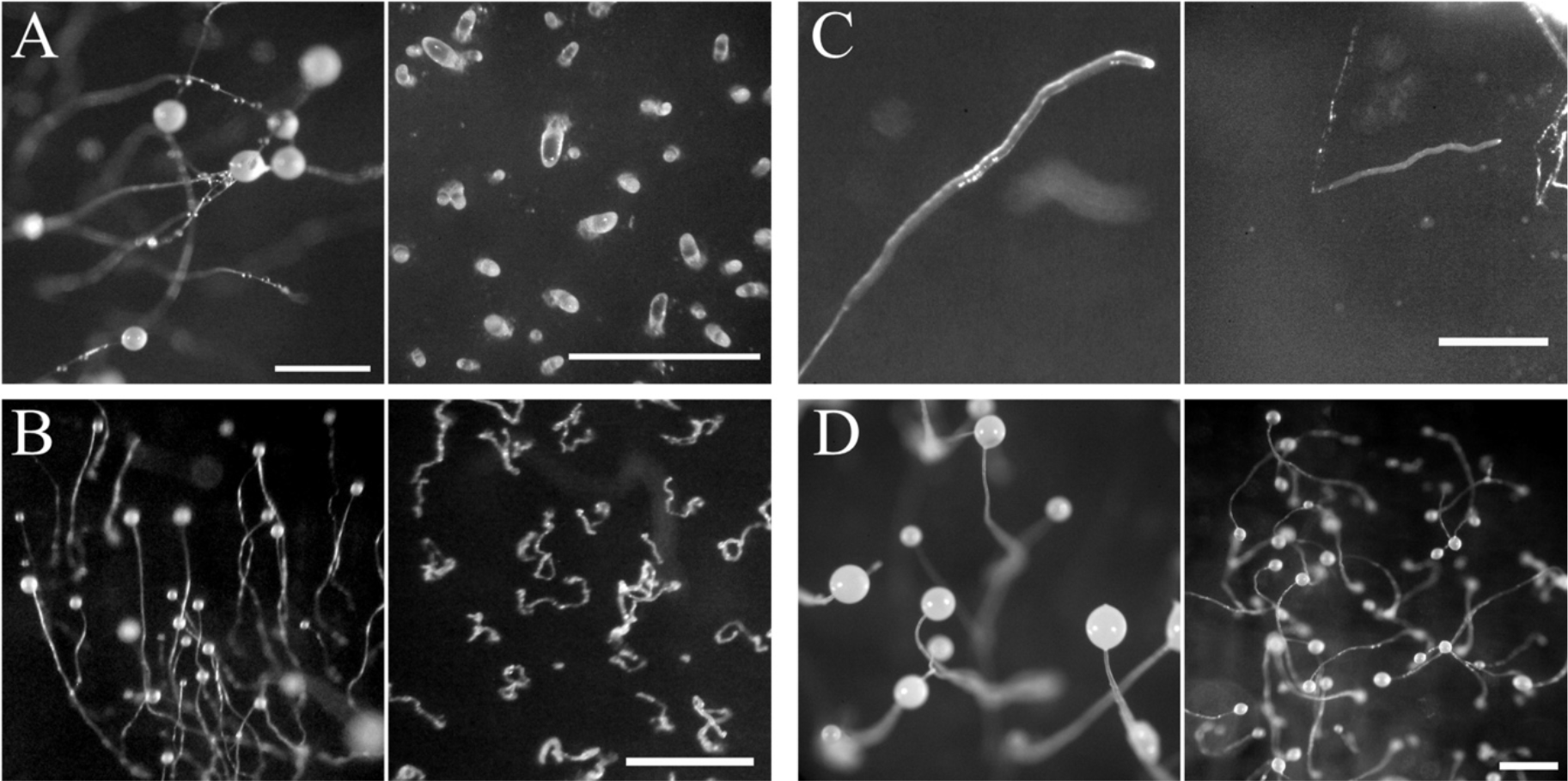
Examples of normal and aberrant structures formed at low and high temperatures (A, B), and normal structures formed at extreme temperatures (C, D). (A) *D. firmibasis* I2Ab. Normal fruiting bodies formed at 16°C (left) and aberrant structures formed at 4°C (right), (B) *D. fasciculatum* (*Cavendaria fasciculatum*) Smok. Normal fruiting bodies formed at 22°C (left) and aberrant structures formed at 28°C (right), (C) *D. polycephalum* (*C. polycephalum*) 1JND1. Normal migrating slugs formed at 16°C (left) and 34°C (right), (D) *D. septentrionalis* (*D. septentrionale*) AK2. Normal fruiting bodies formed at 16°C (left) and 0°C (right). Photographs on the right hand side of each panel were taken after 10 days (A), 6 days (B), 8 days (C), and 13 days (D) of development on non-nutrient agar at the indicated temperatures, after which there were no noticeable morphology changes over the period of at least one month. Scale bar, 1mm.

Fig 3 summarises the results of 36 dictyostelid species (S5 Table). In each graph, the growth rate relative to the maximal growth rate of that species is plotted against temperature (S2 Fig for growth rates of other strains/isolates), while the final form of development is diagrammatically shown in the next column. All species grew and formed fruiting bodies over a wide range of temperatures. This tolerance range was around 20 degrees, but some species showed considerably wider or narrower ranges. For example, *D. mucoroides* (strain no.7 = *D. clavatum*, see below) grew and developed over a range of 30 degrees, while *D. discoideum* (strain NC4) showed much narrower tolerance. The tolerance ranges of growth and development were mostly in good agreement (Fig 3, S3 Fig). It has been thought that the optimal temperature for growth in dictyostelids generally range between 20°C and 25°C [14], with some exceptions such as *D. polycephalum* (*C. polycephalum*) showing optimal growth at 27°C or higher [33] and *D. septentrionalis* (*D. septentrionale*) showing optimal growth at 15-17°C [17]. Over half of the species investigated in this study grew maximally and formed apparently normal fruiting bodies at 28°C or above. They may be called “thermophilic” species. Among the most thermophilic was one strain of *D. polycephalum* (*C. polycephalum*), which grew and formed normal slugs at 34°C (Fig 2). It has been reported that *P. pallidum* (*H. pallidum*) strain “Salvador” could grow and form normal fruiting bodies at 37°C [13],[14], but no species investigated in this study grew or developed at 37°C. On the other hand, some species grew and formed normal fruiting bodies at 4°C (“cold-tolerant” species hereafter). Surprisingly, some of those were still able to grow and aggregate even at 0°C. Among them, the most cold-tolerant species, *D. septentrionalis* (*D. septentrionale*) and *D. brefeldianum*, formed normal fruiting bodies, albeit very slowly, at this temperature (Fig 2). To our knowledge, this is the first report of growth and development of dictyostelids at 0°C.

**Fig 3.**
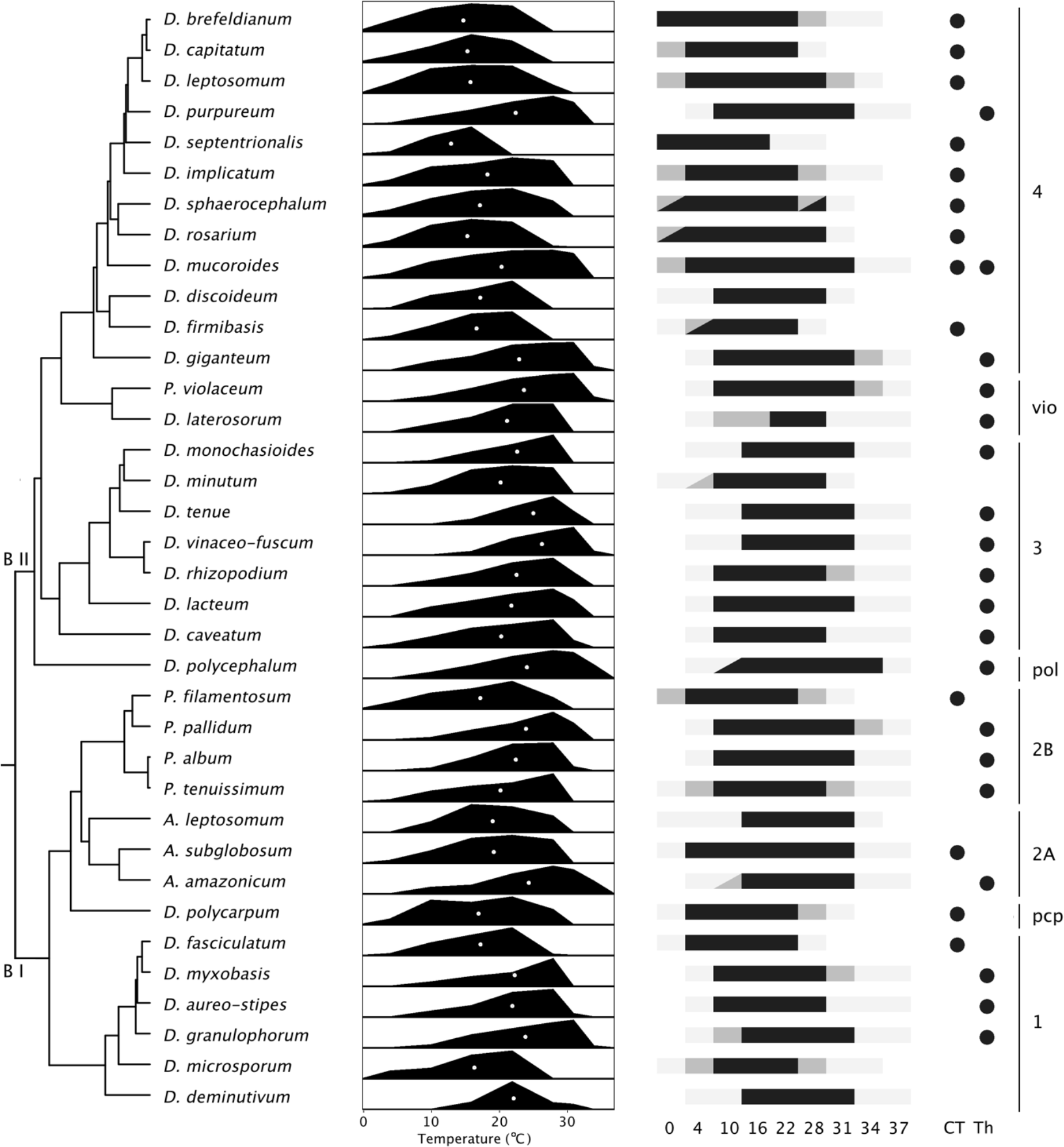
Summary of the temperature dependency of growth and development in 36 dictyostelid species. From left to right, phylogenetic tree, relative growth rate plotted against temperature, final form of development (black: fruiting body, dark grey: aberrant multicellular structure or incomplete development, light grey: only aggregation), thermal characteristics (CT: cold-tolerant, Th: thermophilic), and clade. The white dots in the graph indicate the mean growth temperature (first-order moment of the growth-temperature curve). For development, if the final form observed on non-nutrient agar plate differed from that on LP agar plate, they are shown above and below the diagonal line, respectively. Results of additional isolates of 14 species are shown in S2 Fig.

### Thermal characteristics and habitat temperature

The global-scale geographical distribution of dictyostelids has been studied by Cavender and others (Table 1). Comparing the above results (Fig 3) with the distribution pattern, one can see a clear correlation between the thermal characteristics of a species and their geographical distribution; of the 8 tropical species examined in this study, all were thermophilic, and 8 of the 11 subarctic species were cold-tolerant. For further analysis, however, simply correlating the thermal characteristics with the known geographical distribution by species name will be insufficient because of possible within-species variation and the broad distribution of the many well-known species [11]. Fig 4 shows the correlation between the thermal characteristics of the individual isolates and the annual average temperature of their collection site. The correlation with the “optimal growth temperature” estimated from the growth-temperature curve (see Materials and methods) was weak and not significant (Fig 4A). As another measure for temperature preference, we take the first-order moment of the growth-temperature curve, which is the “centre of mass” of the growth-temperature relationship and may be called the “mean growth temperature” (shown in white dots in the graphs of Fig 3). There was a statistically significant relationship between them, but the scatter of the points is still large (Fig 4B). The scatter, when viewed in the vertical direction, indicates that many of them originated from areas far apart from their favoured habitats as far as average temperature is concerned. When viewed in the horizontal direction, the scatter indicates that species with diverse thermal traits may inhabit areas with similar temperature conditions. For example, of the 11 isolates of 10 species (encompassing the 8 dictyostelid clades) isolated from soils collected at a single site (open symbols in Fig 4), 6 were thermophilic and 4 were cold-tolerant, with their mean temperatures for growth ranging from 14.1°C (*D. brefeldianum*, one of the most cold-tolerant species), to 24.2°C (the most thermophilic isolate of *D. purpureum,* 1V2Ba strain, examined in this study, S2 Fig.). These results indicate that they may inhabit areas with diverse temperature environments without adaptation to local conditions.

**Table 1.** Dictyostelid species characteristic of tropical and subarctic regions taken from literature and their phylogenetic position.

| Tropical species | Group | Subarctic species | Group |
| --- | --- | --- | --- |
| *D. purpureum* | 4 | *D. brefeldianum* | 4 |
| *D. macrocephalum* | 4 | *D. capitatum* | 4 |
| *D. laterosorum* | Pvio | *D. leptosomum* | 4 |
| *D. monochasioides* | 3 | *D. aureocephalum* | 4 |
| *D. tenue* | 3 | *D. septentrionalis* | 4 |
| *D. rhizopodium* | 3 | *D. implicatum* | 4 |
| *D. vinaceo-fuscum* | 3 | *D. crassicaule* | 4 |
| *D. coeruleo-stipes* | 3 | *D. sphaerocephalum* | 4 |
| *D. lavandulum* | 3 | *D. discoideum* | 4 |
| *D. polycephalum* | Dpol | *D. minutum* | 3 |
| *P. pseudo-candidum* | 2B | *P. filamentosum* | 2B |
| *D. granulophorum* | 1 | *P. candidum* | 2B |
|  |  | *D. fasciculatum* | 1 |
|  |  | *D. aureo-stipes* var. *helveticum* | 1 |
|  |  | *D. microsporum* | 1 |
|  |  | *D. parvisporum* | 1 |

The list is based on the previous reports [11],[34]–[36]. Phylogenetic groups are based on the current classification, Schilde., et al (2019) [9].

**Fig 4.**
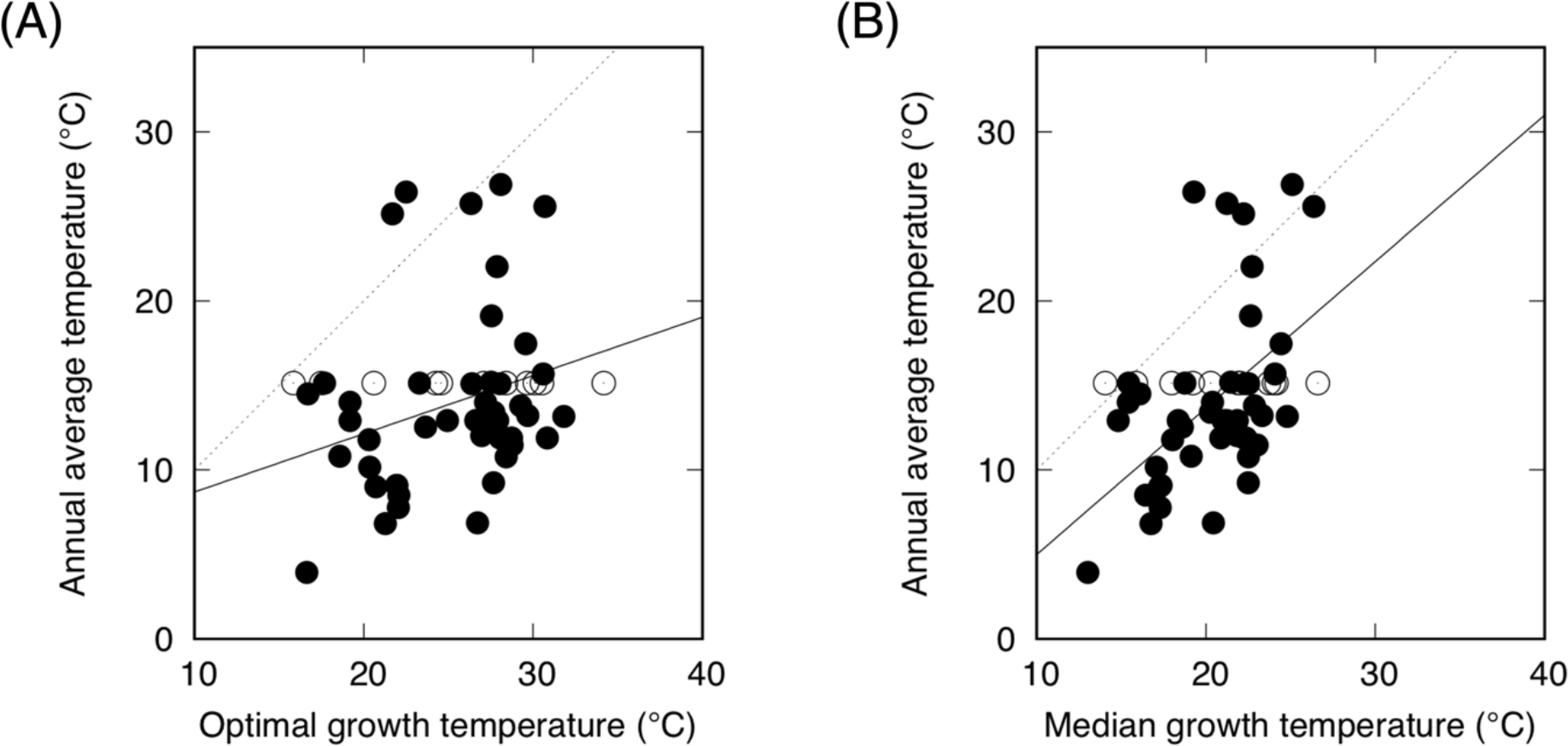
Relationship between (A) optimal and (B) mean growth temperature of all strains/isolates and the annual average temperature of their collection site. Open symbols are those obtained from a single site to illustrate the range of thermal characteristics of the species occurring in the same location (Fig 6 for detail). Solid lines are the linear regression lines for data points shown in closed symbols. Correlation coefficient and 95% C.I. are 0.269 and 0.027–0.578, respectively, for (A), 0.500 and 0.242–0.692 for (B). Dashed lines: the line of equality between annual average temperature and optimal/mean growth temperature.

To gain insight into how their geographical distribution is related with seasonal temperature changes, we focused on the relationship between the tolerance limits of individual isolates and the monthly average temperatures of their collection site. As measures of the lower and upper temperature limits for growth, we use the temperatures corresponding to about 10% maximal growth (LL10 and UL10, see Material and Methods: S5 Table). Fig 5A shows the relationship between the monthly average temperature of the coldest month at the collection sites and LL10 for the individual isolates examined in this study. Fig 5B shows the relationship between the monthly average temperature of the 6^th^ warmest month and LL10. The zone below the line of equality (diagonal) line represents the zone of <10% maximal growth because of low temperature. The monthly average temperature was well below LL10 during cold months in many isolates (37 of the 56 isolates in the coldest month, Fig 5A), and it was only in the 6^th^ warmest month that the monthly average temperature exceeded LL10 of all isolates (Fig 5B). In similar analyses with the temperature range giving ≥50% maximum growth, the monthly average temperature was lower than the lower limit of the range of ≥50% maximal growth (LL50) during cold months in many isolates, and it was only in the second warmest month that the monthly average temperature became higher than LL50 of all isolates (Figs 5C and 5D). For the upper limit, UL10 was higher than the monthly average temperature of the warmest month in most isolates analysed. These results mean that dictyostelids can inhabit areas where the monthly average temperature remains below LL10 for up to half year and below LL50 for up to 10 months. This may not be surprising because spores and cysts survive temperature conditions outside the tolerance limit, but the results demonstrate that they manage adverse conditions even in the presence of other species more adapted to the location. When looked at individually, maximal growth of the individual isolates will be attained at different times of the year depending on their thermal characteristics and the temperature conditions of the collection site. In the example shown above of the 10 species isolated at a single site in the temperate region, many will show maximal growth during the summer months whereas maximal growth of the cold-tolerant species is predicted to occur in spring and autumn (Fig 6).

**Fig 5.**
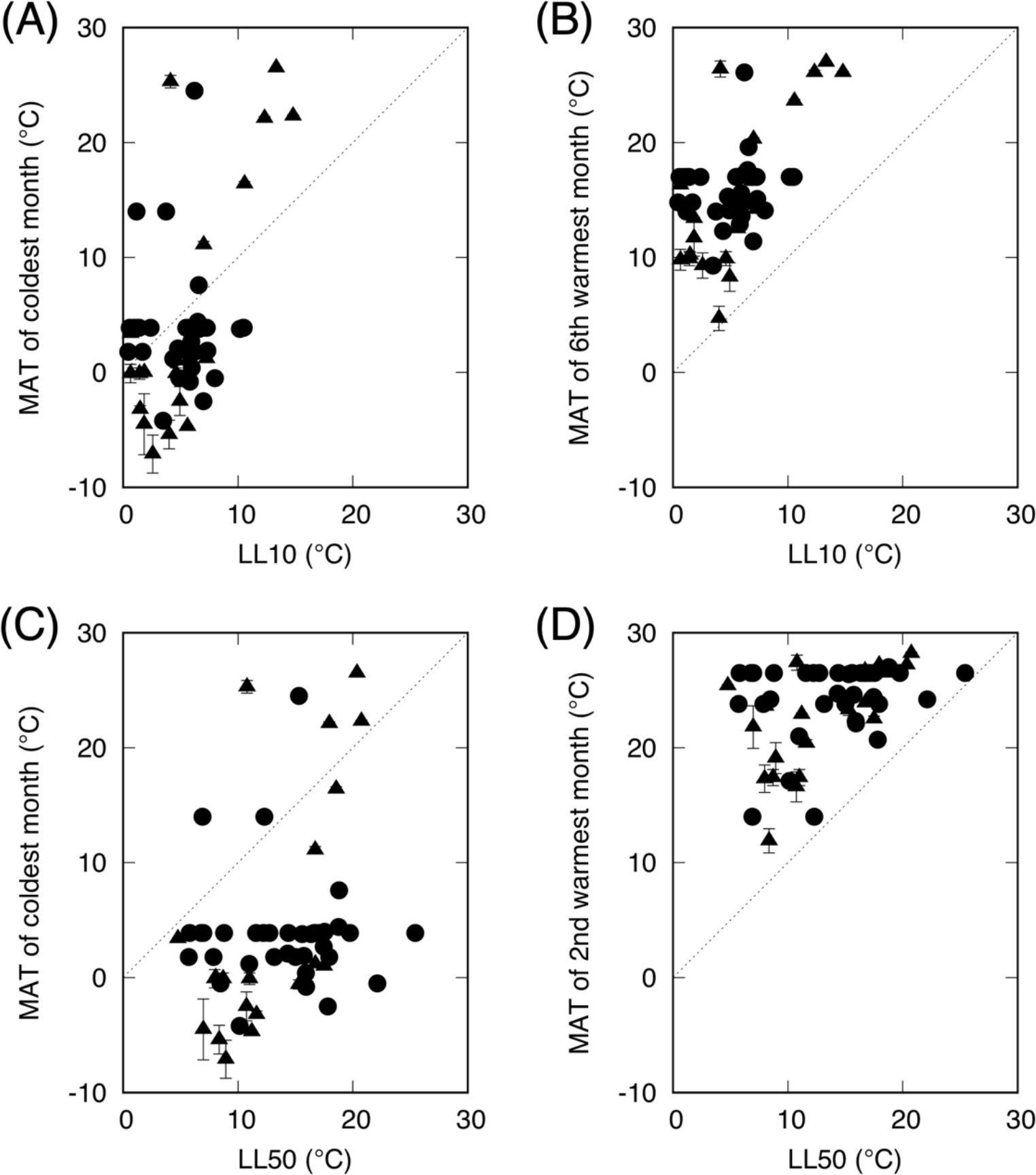
Relationship between the upper or lower temperature limit for growth and the monthly average temperature (MAT) of a particular month at their site of origin. Circles: strains/isolates of which the precise location of collection site is known, triangles: those of which the precise location of collection site could only be obtained as an area extending over 0.5 min in longitude/latitude. Error bar indicates the range between the maximum and minimum average temperatures within the estimated area. Each triangle symbol is plotted at the mean of the temperatures in that area. (A) MAT of the coldest month of the year at the site of origin plotted against the lower temperature limit of the range of ≥10% maximal growth (LL10). (B) MAT of the sixth warmest month of the year at the site of origin plotted against LL10. (C) MAT of the coldest month of the year at the site of origin plotted against the lower limit of the temperature range giving ≥50% maximum growth (LL50). (D) MAT of the second warmest month of the year at the site of origin plotted against LL50. The straight lines in (A) and (B) are MAT = LL10, those in (C) and (D) are MAT = LL50.

**Fig 6.**
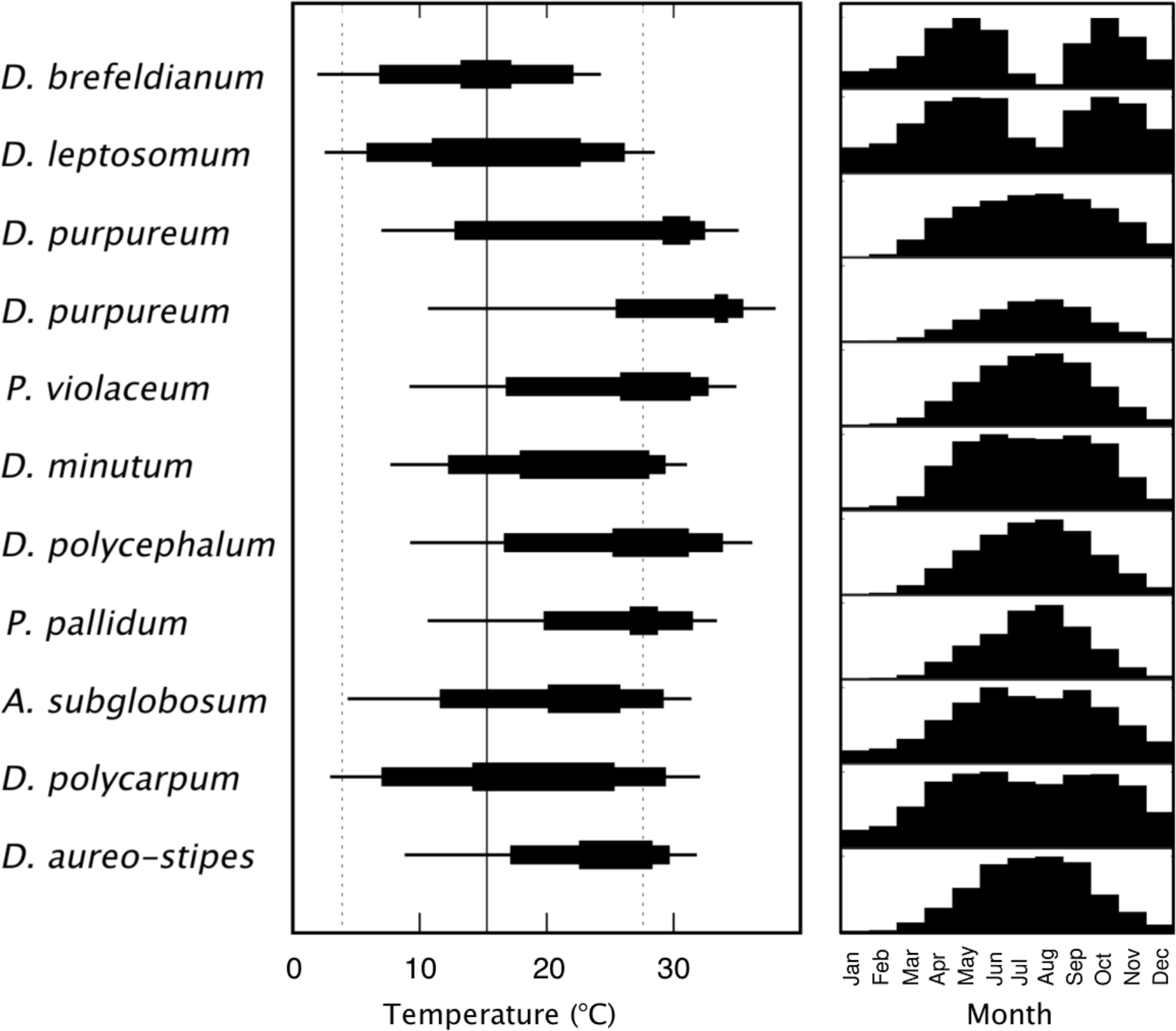
Relative growth rate of the 10 species (11 isolates) collected at a single site. They were isolated from several soil samples collected in a wooded area (1.5 ha) of the botanical garden of Kyoto University Graduate School of Science in May (2011 to 2015). Left panel: temperature effects on the relative growth rate (from Fig 3 and S2 Fig) shown diagrammatically. Thin horizontal lines show the temperature range between LL10 and UL10 (ca. 10% maximal growth), medium thickness lines the range between LL50 and UL50 (50% maximal growth), and thick lines the range of over 90% maximal growth rate. The solid vertical line indicates the annual average temperature of the site. The dashed lines are the monthly average temperatures of the coldest month (January) and warmest month (August). Right panel: average relative growth rate at each month deduced from the temperature-dependent relative growth rate of each isolate and the monthly average temperature of the site.

### Phylogenetic association of thermal characteristics

Combining the list of dictyostelids distribution with the current knowledge of the dictyostelid phylogeny [6]–[9], one can see clear correlation between distribution and phylogeny (Table 1). Notably, about half the species reported as tropical species [11],[34] belong to group 3, and the great majority of group 3 species are restricted to the tropic to sub-tropic zones [14]. In contrast, about half the species recorded in the sub-arctic regions belong to group 4 [11],[34]–[36]. Recently, it has suggested that survival activities of spores against low temperature are prominent in group 4 species which were frequently isolated from the cold climate zones [37]. It may be inferred that the phylogenetically constrained thermal tolerance of individual species influences their geographical distribution.

It can be seen in Fig 3 that the temperature ranges for growth and development of many group 4 species are lower than most of the other groups, suggesting a possible correlation with phylogeny. Over 70% of the group 4 species investigated were cold-tolerant, whereas all but two of the group 3, clade 2B and two group-intermediate clades (violaceum and polycephalum) were thermophilic. The mean growth temperature of the group 4 and clade polycarpum were significantly lower than that of all other clades except clade 2A (*P* < 0.05 to *P* < 0.005, unequal variances t-test). The difference in the lower limit of growth temperature was even clearer, while the difference in the upper limit was relatively small (Table 2). These results indicate that the thermal characteristics of dictyostelids are influenced by phylogeny to some degree. This was supported by the widely-used measure of phylogenetic signal, Pagel’s λ [26]. In this index, random changes of the traits over evolution give values close to one, whereas diversification under strong selection pressure would give a value close to zero. If calculated for the entire tree of the 36 species, estimates of Pagel’s λ for the mean, lower limit, and upper limit of temperature for growth were 0.58, 0.58, and 0.43, respectively, indicating the presence of phylogenetic signal for the mean and lower limit of temperature for growth (S6 Table). Branch II, which consists of groups 3 and 4 and two minor clades (22 species) gave much stronger signal (0.75, 0.76, 0.57) for the mean, lower limit, and upper limit, respectively. Two other measures of phylogenetic signal, Blomberg’s K [27] and Moran’s I [28] gave similar results (S6 Table).

**Table 2.** Number of cold-tolerant and thermophilic species, and mean, lower limit, and upper limit temperature for growth.

| Clade | number of strains |  |  | growth temperature (°C) |  |  |  |  |  |  |  |  |
| --- | --- | --- | --- | --- | --- | --- | --- | --- | --- | --- | --- | --- |
|  | total | cold-tol-<br>erant | thermophilic | median | *a* | *b* | lower limit | *a* | *b* | upper limit | *a* | *b* |
| group 4 | 19 | 13 | 5 | 18.03 ± 3.45 | — |  | 4.15 ± 2.57 | - |  | 29.42 ± 4.98 | — |  |
| violaceum | 4 | 0 | 4 | 22.70 ± 1.09 | *** | *** | 8.72 ± 0.34 | *** | *** | 33.05 ± 1.87 | * |  |
| group 3 | 7 | 0 | 6 | 22.77 ± 2.28 | *** | *** | 10.01 ± 3.77 | *** | *** | 32.69 ± 1.17 | ** |  |
| polycephalum | 4 | 0 | 4 | 21.96 ± 1.48 | *** | ** | 7.92 ± 0.88 | *** | *** | 33.33 ± 2.14 | * |  |
| clade 2b | 6 | 1 | 5 | 21.38 ± 2.33 | ** | ** | 7.58 ± 3.08 | * | ** | 32.58 ± 1.74 | * |  |
| clade 2a | 4 | 1 | 1 | 20.52 ± 2.61 |  |  | 6.80 ± 2.13 | — | * | 32.30 ± 2.64 |  |  |
| polycarpum | 2 | 2 | 0 | 17.52 ± 0.64 |  | — | 3.10 ± 0.28 | — | — | 31.25 ± 1.20 |  | — |
| group 1 | 8 | 2 | 5 | 20.73 ± 2.70 | * | ** | 8.00 ± 3.57 | ** | *** | 30.82 ± 3.38 |  |  |

*a*: statistical significance of the difference between the average of that group and that of group 4. *b*: statistical significance of the difference between the average of that group and that of the polycarpum complex. \* $P < 0.05$ , \*\* $P < 0.01$ , \*\*\* $P < 0.0005$ .

We analysed the correlation between trait divergence and phylogenetic distance for all isolates examined in this study. Fig 7 shows the difference in the mean growth temperature between two isolates plotted against square root of the phylogenetic distance (branch length) between them for all combinations of the isolates. Differences within each group are shown in colour and those between groups in grey. In group 4 (purple symbols), the maximum divergence increases approximately linearly as a function of square root distance, indicating that the diversification of mean growth temperature is largely a random process as in Brownian motion [38]. Notably, the rate of divergence in group 4 is higher than the other groups, and group 4 in itself exhibits the full range of dictyostelid diversity. Ancestral state reconstruction using a maximum-likelihood method based on a model of trait evolution under Brownian motion suggests the presence of a strong trend towards cold-tolerance in group 4 whereas the acquisition of tolerance to high temperature is scattered over the entire tree (Fig 8).

**Fig 7.**
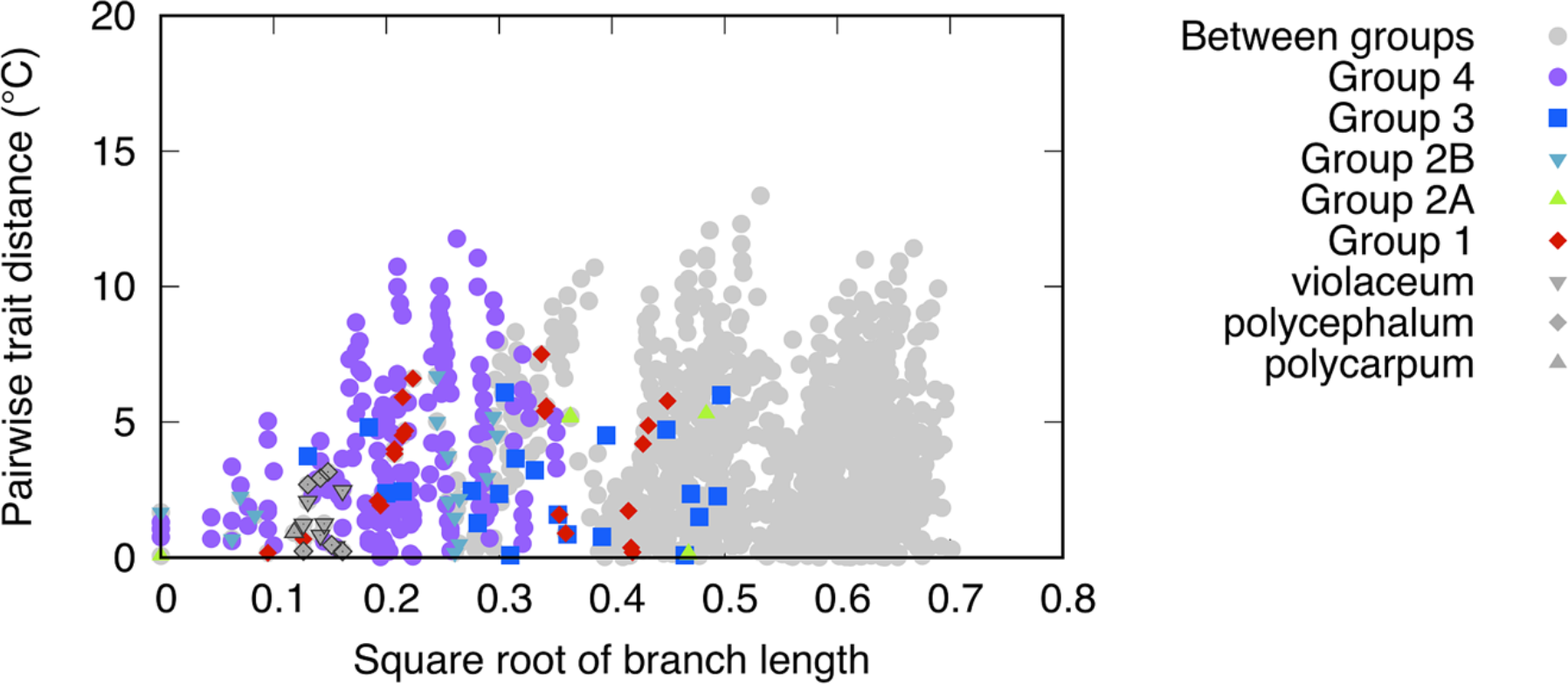
Pairwise difference in the mean growth temperature plotted against square root of the phylogenetic distance for all combinations of the 54 isolates examined. The distance is the sum of the branch lengths on a tree constructed by embedding local trees into the dictyostelid phylogenetic tree [7].

**Fig 8.**
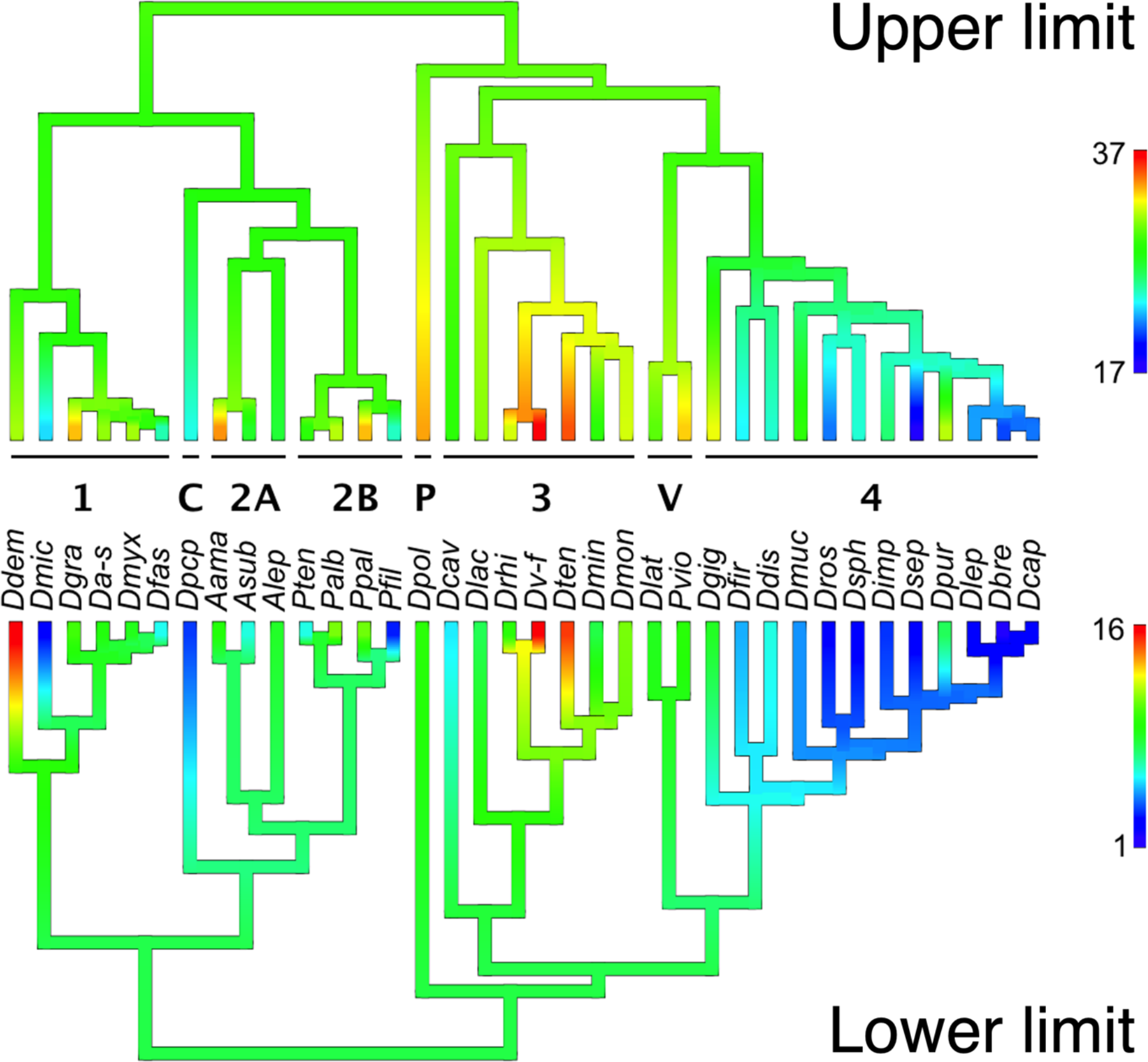
Estimation of ancestral upper and lower temperature limits for growth (reconstructed using a maximum-likelihood-based method). Top: upper limit, bottom: lower limit. Species names are abbreviated (see S1 Table). The numbers of the major groups and abbreviations for the minor clades are shown above the species names. C: polycarpum, P: polycephalum, V: violaceum. Estimates and 95% confidence intervals for the lower limit of the root of these clades are as follows (in degrees Celcius). Group 1: 9.02 (4.15 – 13.89), group 2A: 7.25 (2.28 – 12.2), group 2B: 7.28 (3.42 – 11.14), group 3: 7.82 (2.16– 13.49), group 4: 5.70 (1.45 – 9.95), polycarpum: 3.29 (0.92 – 5.65), polycephalum 8.05 (5.40 – 10.72), violaceum: 8.23 (3.40 – 13.05).

## Discussion

### Thermophicility and cold-tolerance of dictyostelids

Over half of the species investigated grew and formed fruiting bodies maximally at ≥ 28°C, which is higher than the temperatures generally considered optimal for most species [14]. On the other hand, we found that some “cold-tolerant” species can grow and form apparently normal fruiting bodies at 4°C, some even at 0°C. Low temperatures generally inhibit cell activities such as cytokinesis and cell differentiation, and this has been shown also for *D. discoideum* as well [18],[39]. The cold-tolerant dictyostelids may have mechanisms for maintaining their activities at low temperature, such as stabilisation of the cytoskeleton and alteration of the membrane lipid composition at low temperature, as known in other eukaryotes [40],[41].

### Thermal characteristics and habitat temperature

Literature survey indicates that dictyostelid species collected from the sub-arctic and tropical regions show better growth and development at correspondingly low and high temperatures [14]. On the other hand, the geographical distributions of common species are very broad [11],[34]. Our results demonstrate that many species can inhabit areas with thermal conditions far removed from their optimal, suggesting that their broad distribution is not the result of adaptation to local temperature environments but rather due to their ability to proliferate under a wide range of temperature conditions [42],[43]. The broad distributions may also be due to variation within the species. The presence of ecotypes of *D. discoideum* differing in temperature tolerance has been suggested [44]. However, independent isolates assigned to the same species but significantly differing in thermal characteristics may differ at the species level. For instance, the subtropical and temperate forms of *D. purpureum* differ in many aspects, and are most likely of different species [45],[46]. The two most thermophilic isolates of *D. purpureum* (F1LF, 1V2Ba) were very similar from morphological and molecular viewpoints to the subtropical form (S2 Fig). It was also noted in the present study that the genetic distances between isolates of the same species with distinct thermal characteristics tended to be large (S2 Fig).

The correlation between the thermal characteristics of individual isolates and the monthly temperature data of their collection sites indicates that dictyostelid slime moulds live in areas where temperature conditions permit proliferation during a limited period of the year; among the isolates examined, none originated from areas where the monthly average temperature was below the lower temperature limit for growth (defined in this study as the temperature giving 10% growth rate relative to the optimal growth) over half year (Fig 5). In other words, at least half year of minimal growth period would be needed for a species to establish. In the wild, different species may coexist in a very small patch of soil [47],[48] and compete each other for food source and niches [49], so the one with thermal characteristics more suited to the temperature conditions of that region would predominate if other conditions are equally favourable, as was demonstrated in laboratory experiments [13]. Small differences in the thermal characteristics among competing species could thus lead to their large-scale, graded distributions along the climatic gradients, such as the latitudinal and altitudinal distribution patterns documented in earlier studies [34],[50].

Species showing wide thermal tolerance will be able to expand their distribution to broader climate conditions. Such species will remain active for longer periods of the year than those with narrow thermal tolerance, and may also be more adaptive to environments where annual and diurnal temperature changes are large. It is also conceivable that the difference in temperature preference between species might cause seasonal variation of the population structure. Monthly relative growth rates deduced from the temperature dependence of growth and the climatic data predict distinct seasonal growth patterns depending on the thermal characteristics of the species (Fig 6). Such a case can be found in a report in which *P. pallidum* (*H. pallidum*), a thermophilic species, dominated in summer while *D. mucoroides*, a cold-tolerant species, were the most abundant during winter in a temperate forest soil in Japan [51]. In two other studies conducted in Europe and North America, there is no clear relationship with the thermal characteristics in the changes of species composition [52],[53]. Presumably, the effects of periodically changing temperature are, unlike the climate effects over a geological time scale, not cumulative, and could be readily overridden by other physical and biological factors.

## Phylogenetic association of thermal characteristics

The distribution of the thermal characteristics over the dictyostelid phylogenetic tree suggested a link between the traits and phylogeny (Table 2). This relationship was substantiated by the clear correlation between trait differences and phylogenetic distance (Fig 7) and further supported by the estimates of widely-used measures of phylogenetic signal (S6 Table). The ancestral trait reconstruction (Fig 8) suggests that cold-tolerance evolved several times independently throughout the dictyostelid phylogeny in all of the major clades and one minor clade, and that the one evolved in group 4 diversified and expanded their distribution extensively. This agrees well with the recent findings by Lawal et al. that spores of group 4 species show enhanced cold-resistance during dormancy due to thicker spore walls than other groups [37]. For the evolution of thermophilicity, on the other hand, the predominance of thermophilic species in major taxonomic groups is consistent with the previous suggestion by Cavender [34] that most speciation occurred in the tropics.

## Phylogeny and distribution

A clear correlation between phylogeny and distribution can be found in the literature [11],[34]–[37] (Table 1). Many of the species characteristic of the tropics and subtropics belong to group 3, and many isolates from the cold regions are in group 4. Our results indicate strong associations between thermophilicity and group 3 species and between cold-tolerance and group 4 species (Table 2). They further suggest the possibility that the thermal characteristics of these species permitted their spreading to areas with climatic conditions in which they were capable of proliferation. The fruiting body of group 4 species is generally large and robust with a thick stalk [7],[14]. This, together with cold-tolerance, would be advantageous in harsh conditions of the subarctic environments [34],[37]. The phylogenetic association of traits and geographical distribution could also be explained if the spreading of the organism is slow, such that its descendants would diverge under similar environmental pressure. However, this is unlikely to be the main cause for the dictyostelid slime mould, as its spreading by birds and large animals is most likely very efficient [48],[54]. In fact the same or closely-related species have been found in very distant regions, often on different continents, of similar climatic conditions [11],[55]. Our phylogenetic analysis (S6 Table) also argues against rapid adaptive responses to be the main cause of the correlation between phylogeny and distribution in dictyostelids. From the relationship between the maximum divergence of the mean growth temperature and the phylogenetic distance (Fig 7), together with the results of recent dating studies [37],[56], it may be argued that changes in temperature preference would have proceeded gradually.

## Conclusions

This study substantially increased the number of species with quantitative data on thermal traits of dictyostelids. It basically confirmed the results of earlier studies which were based on a single or phylogenetically biased selection of species. The new data obtained from the species chosen to cover the breadth of the known dictyostelid phylogeny enabled us to compute statistical significance of hypotheses on the evolution of thermal traits in the taxonomic groups. The results imply that the molecular basis for thermal adaptation is genetically conserved, and that the distribution is not the outcome of local adaptation. Specifically, it can be argued that the predominance of cold-tolerant species in group 4 was due to inheritance from ancestors that had accumulated genetic alterations enabling growth and development to proceed under cold conditions. The increased resistance of spores in group 4 species [37] would help such cells to survive the harsh conditions.

## Authors’ Contributions

H.H. conceived the study, planned and performed experiments, and analyzed data. K. I. contributed to the collection and identification of wild isolates, performed phylogenetic analysis and additional experiments. Both authors wrote the manuscript.

## Acknowledgements

The authors would like to thank Kaori Tanabe for technical assistance and Vidyanand Nanjundiah for invaluable suggestions. H.H. was supported by the RIKEN JRA program. This work was supported in part by JP20J00751 and JP21K15081 (to H.H.).

## Competing interests

The authors declare no competing interests.

## Notes

### Competing Interest Statement

The authors have declared no competing interest.

