## Supplementary_information for "Influence of temperature on growth and development of dictyostelid slime moulds and its implication on the evolution of cold-tolerance"

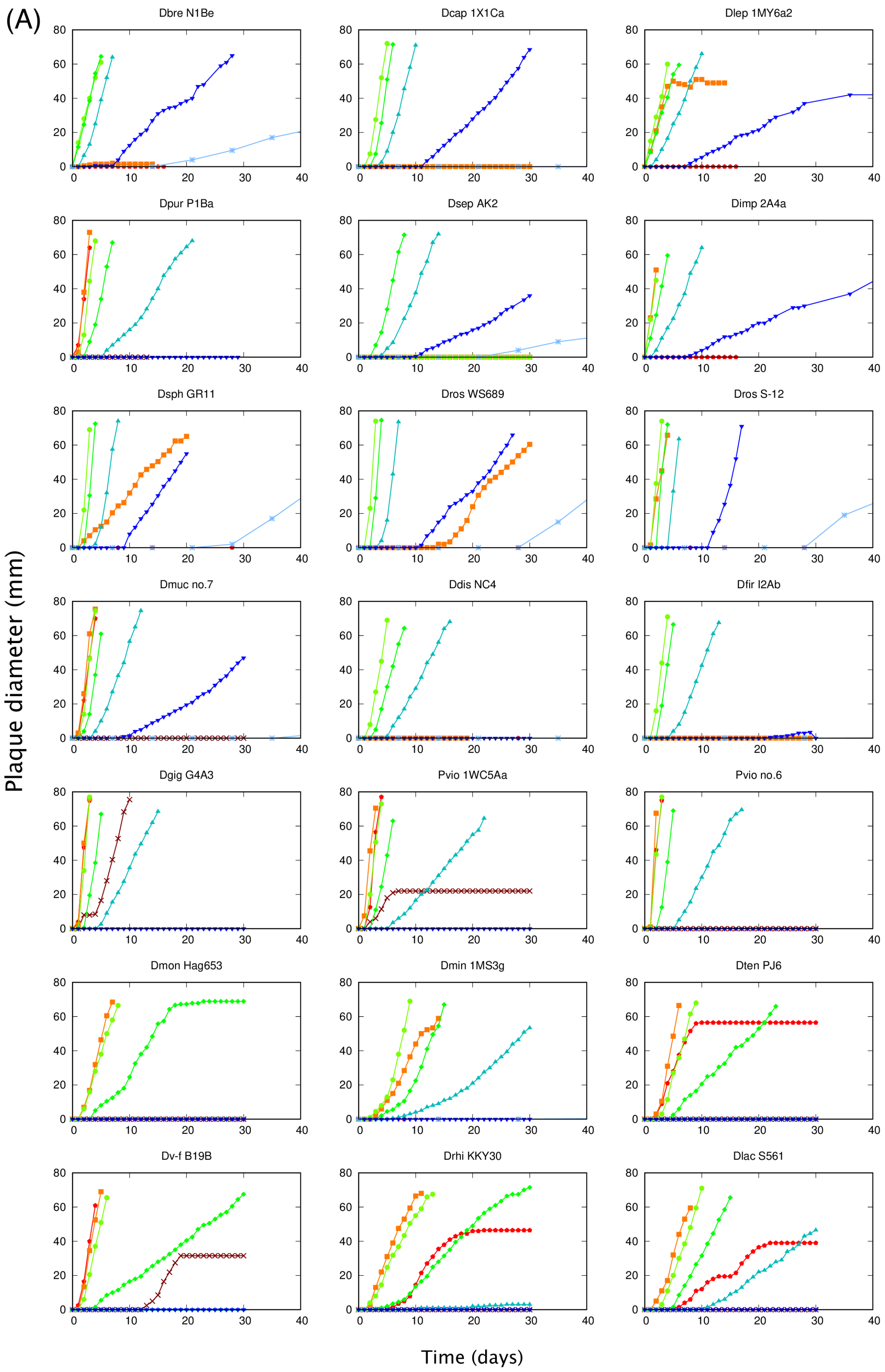
(A)
Dbre N1Be
Dcap 1X1Ca
Dlep 1MY6a2
Dpur P1Ba
Dsep AK2
Dimp 2A4a
Dsph GR11
Dros WS689
Dros S-12
Dmuc no.7
Ddis NC4
Dfir I2Ab
Dgig G4A3
Pvio 1WC5Aa
Pvio no.6
Dmon Hag653
Dmin 1MS3g
Dten PJ6
Dv-f B19B
Drhi KKY30
Dlac S561
Plaque diameter (mm)
Time (days)

### S1 Fig (A) (continued)

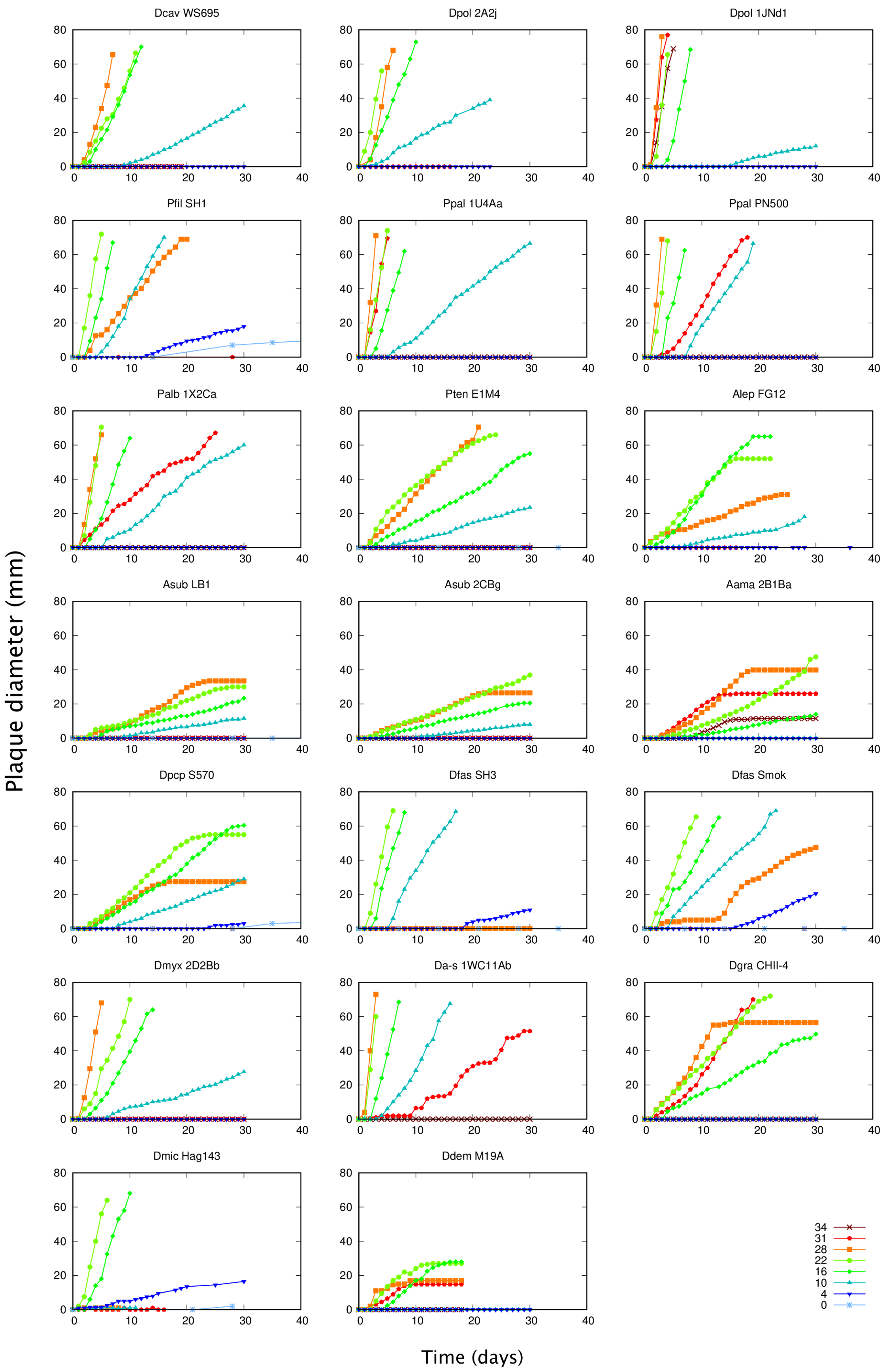

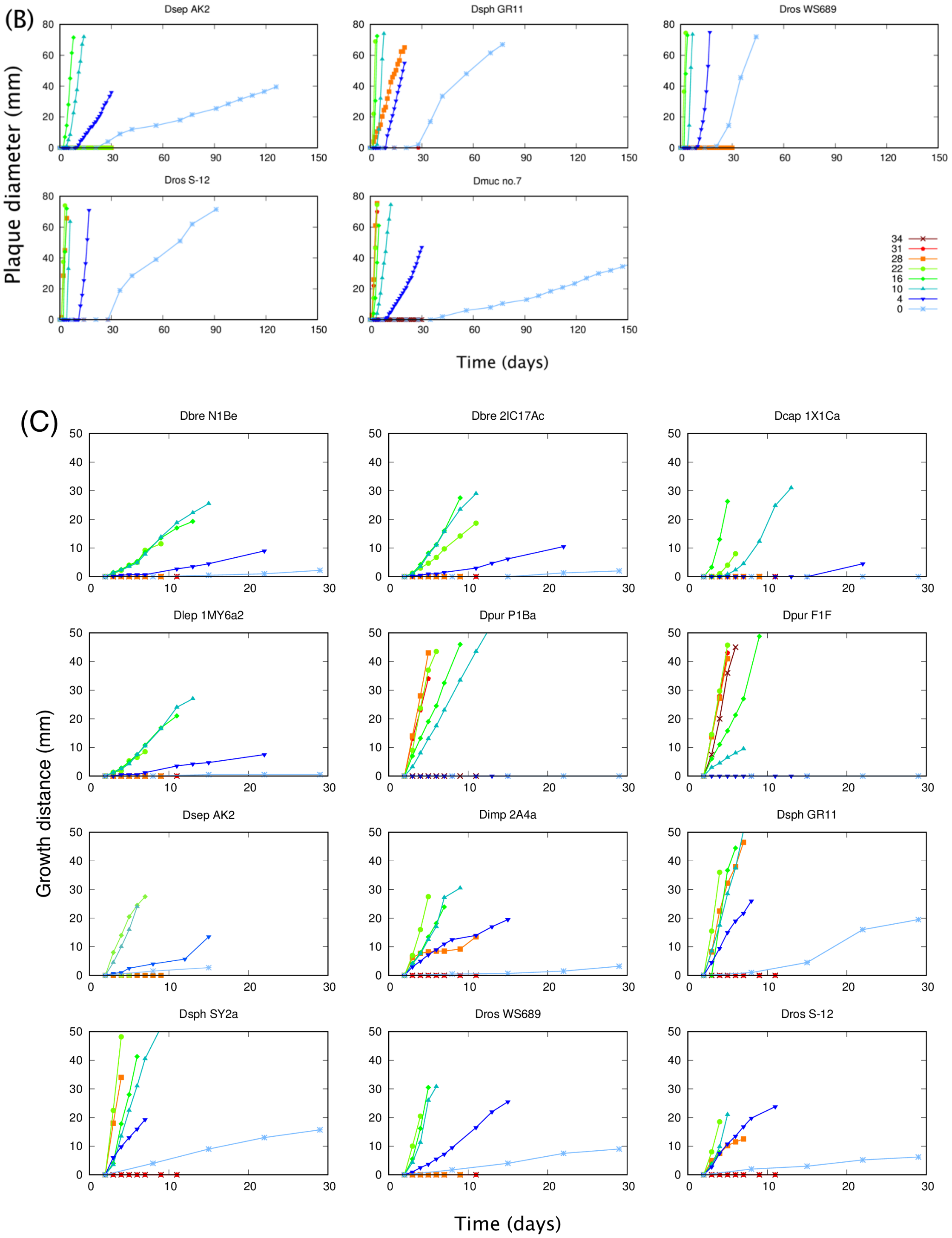

(B)
Plaque diameter (mm)
Dsep AK2
Dsph GR11
Dros WS689
Dros S-12
Dmuc no.7
34
31
28
22
16
10
4
0
Time (days)
(C)
Growth distance (mm)
Dbre N1Be
Dbre 2IC17Ac
Dcap 1X1Ca
Dlep 1MY6a2
Dpur P1Ba
Dpur F1F
Dsep AK2
Dimp 2A4a
Dsph GR11
Dsph SY2a
Dros WS689
Dros S-12
Time (days)

### S1 Fig (C) (continued)

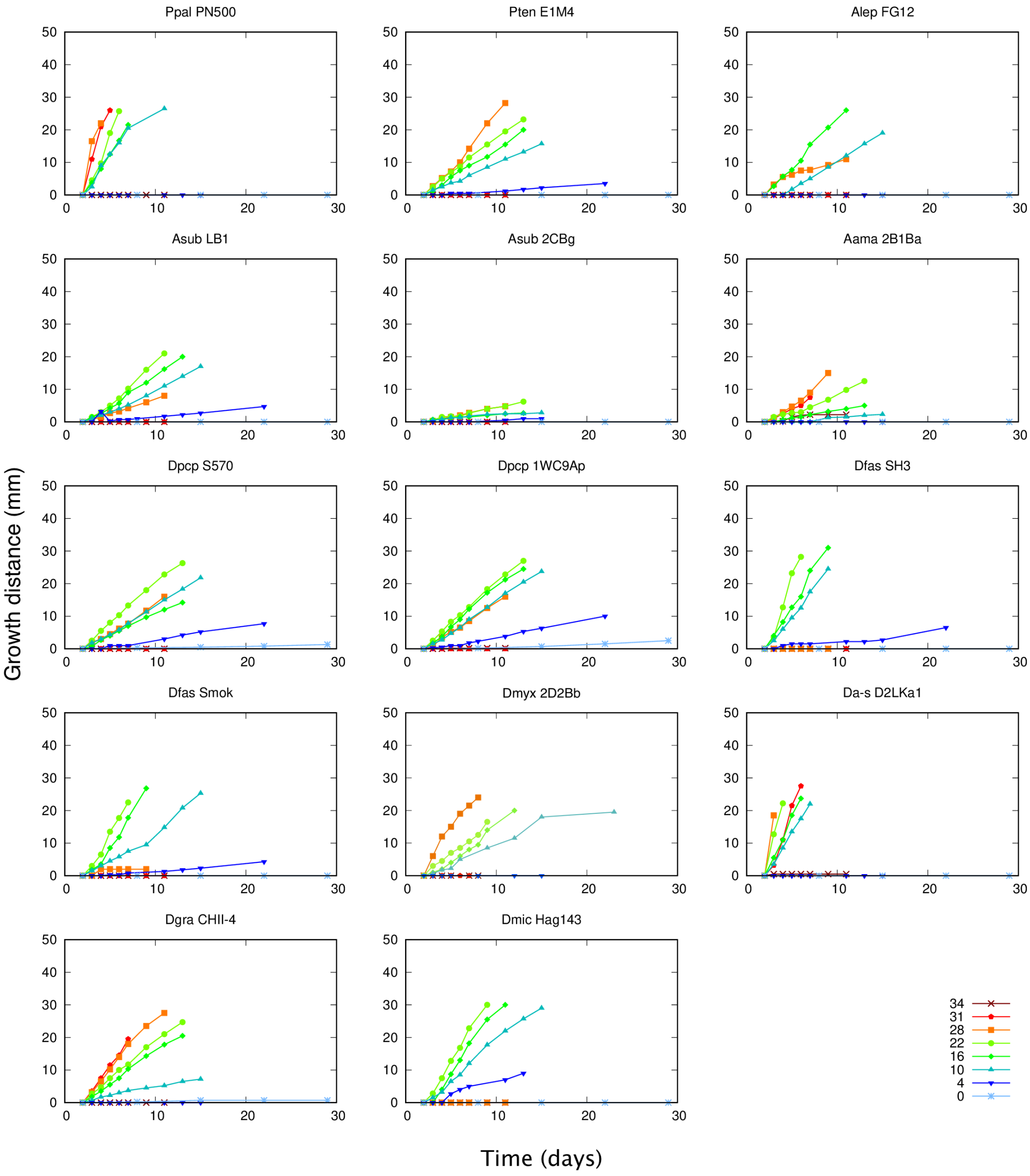


**S1 Fig. Growth curves of all strains/isolates at various temperatures.**

Species names are indicated with the four-letter abbreviation (see S1 Table). (**A)** Growth on a bacterial lawn on nutrient-rich SM medium. (**B)** Data of the 5 isolates (4 species) in (**A)** that showed significant growth at 0˚C are plotted on a larger scale. (**C)** Growth on a bacterial streak on nutrient-poor LP medium.

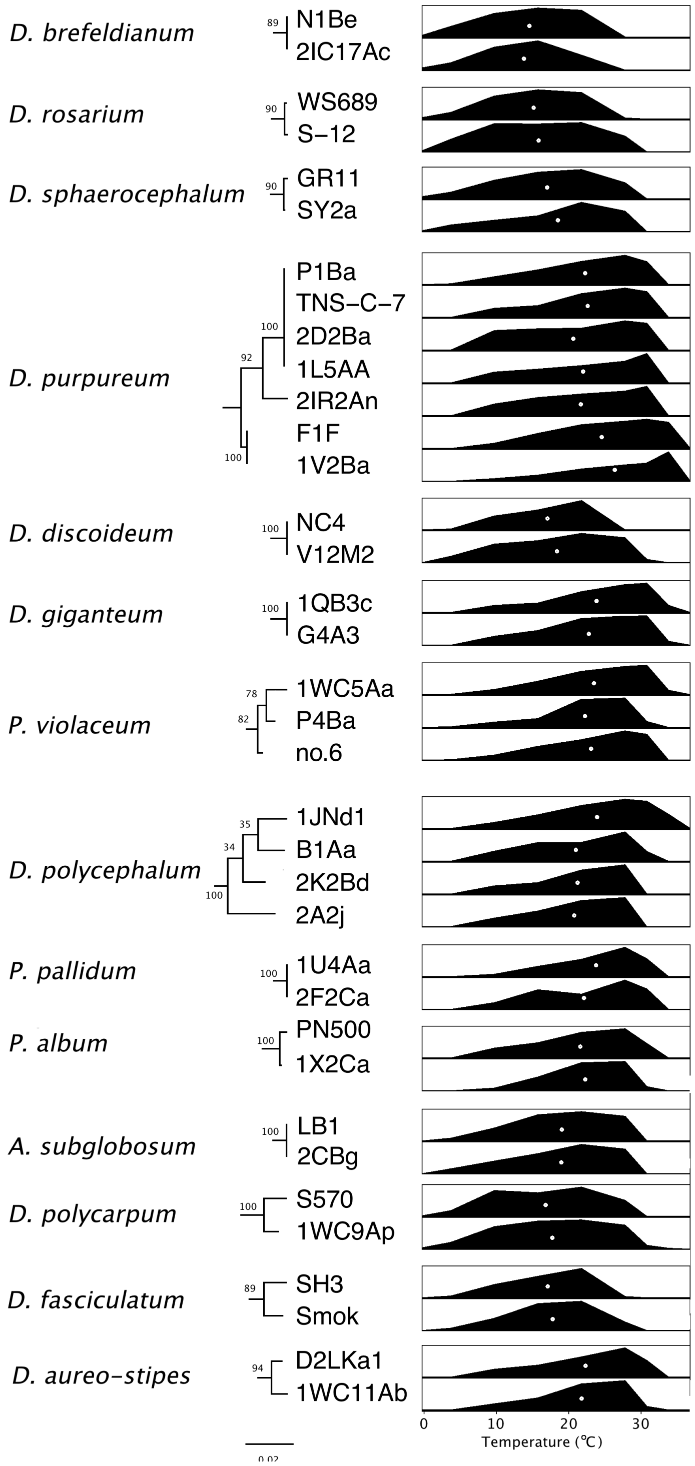


**S2 Fig. Within-species variation of the temperature dependency of growth.**

To the left of the strain/isolate names are partial trees based on SSU rDNA sequences. All trees are shown to scale, as indicated at the bottom of the figure.

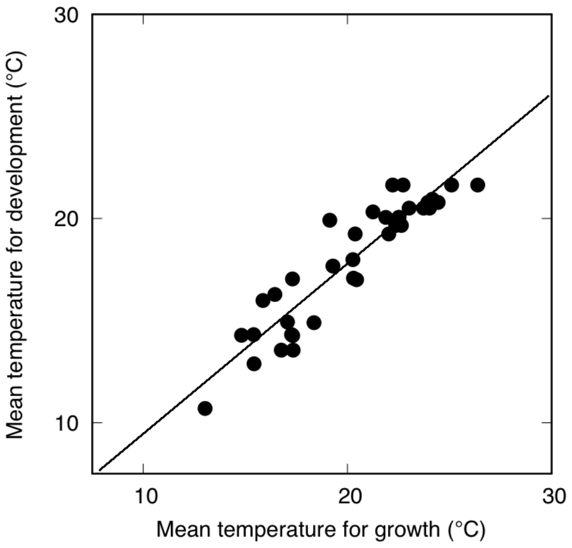


**S3 Fig. Relationship between the temperature ranges for growth and development.** Mean temperature for development of 36 dictyostelid species is plotted against mean temperature for growth. Diagonal line: the line of equality between mean temperature for growth and development.

| clade | Species names[a] | code[b] | strain | clade[c] | Species names[c] |
|---|---|---|---|---|---|
| Group 4 | + *Dictyostelium brefeldianum* | Dbre | N1Be | Genus *Dictyostelium* | *Dictyostelium brefeldianum* |
| | *Dictyostelium brefeldianum* | Dbre | 2IC17Ac | | *Dictyostelium brefeldianum* |
| | + *Dictyostelium capitatum* | Dcap | 1X1Cc | | *Dictyostelium capitatum* |
| | + *Dictyostelium leptosomum* | Dlep | 1MY6a2 | | *Dictyostelium leptosomum* |
| | + *Dictyostelium purpureum* | Dpur | P1Ba | | *Dictyostelium purpureum* |
| | *Dictyostelium purpureum* | Dpur | 1L5Aa | | *Dictyostelium purpureum* |
| | *Dictyostelium purpureum* | Dpur | 2D2Ba | | *Dictyostelium purpureum* |
| | *Dictyostelium purpureum* | Dpur | TNS-C-7 | | *Dictyostelium purpureum* |
| | *Dictyostelium purpureum* | Dpur | 2IR2An | | *Dictyostelium purpureum* |
| | *Dictyostelium purpureum* | Dpur | F1LF | | *Dictyostelium purpureum* |
| | *Dictyostelium purpureum* | Dpur | 1V2Ba | | *Dictyostelium purpureum* |
| | + *Dictyostelium septentrionalis* | Dsep | AK2 | | * *Dictyostelium septentrionale* |
| | + *Dictyostelium implicatum* | Dimp | 2A4a | | *Dictyostelium implicatum* |
| | + *Dictyostelium sphaerocephalum* | Dsph | GR11 | | *Dictyostelium sphaerocephalum* |
| | *Dictyostelium sphaerocephalum* | Dsph | SY2a | | *Dictyostelium sphaerocephalum* |
| | + *Dictyostelium rosarium* | Dros | WS689 | | *Dictyostelium rosarium* |
| | *Dictyostelium rosarium* | Dros | S-12 | | *Dictyostelium rosarium* |
| | + *Dictyostelium mucoroides*[d] | Dmuc | no.7 | | *Dictyostelium mucoroides* |
| | + *Dictyostelium discoideum* | Ddis | NC4 | | *Dictyostelium discoideum* |
| | *Dictyostelium discoideum* | Ddis | V12M2 | | *Dictyostelium discoideum* |
| | + *Dictyostelium firmibasis* | Dfir | I2Ab | | *Dictyostelium firmibasis* |
| | + *Dictyostelium giganteum* | Dgig | G4A3 | | *Dictyostelium giganteum* |
| | *Dictyostelium giganteum* | Dgig | 1QB3c | | *Dictyostelium giganteum* |
| violaceum | + *Polysphondylium violaceum* | Pvio | 1WC5Aa | Genus *Polysphondylium* | *Polysphondylium violaceum* |
| | *Polysphondylium violaceum* | Pvio | P4Ba | | *Polysphondylium violaceum* |
| | *Polysphondylium violaceum* | Pvio | no.6 | | *Polysphondylium violaceum* |
| | + *Dictyostelium laterosorum* | Dlat | AE4 | | * *Polysphondylium laterosorum* |
| Group 3 | + *Dictyostelium monochasioides* | Dmon | Hag653 | Family Raperosteliaceae | * *Raperostelium monochasioides* |
| | + *Dictyostelium minutum* | Dmin | 1MS3g | | * *Raperostelium minutum* |
| | + *Dictyostelium tenue* | Dten | PJ6 | | * *Raperostelium tenue* |
| | + *Dictyostelium vinaceo-fuscum* | Dv-f | B19B | | * *Hagiwaraea vinaceofusca* |
| | + *Dictyostelium rhizopodium* | Drhi | KKY30 | | * *Hagiwaraea rhizopodium* |
| | + *Dictyostelium lacteum* | Dlac | S561 | | * *Raperostelium tenue* |
| | + *Dictyostelium caveatum* | Dcav | WS695 | | * *Speleostelium caveatum* |
| polycephalum | *Dictyostelium polycephalum* | Dpol | 2K2Bd | Genus *Coremiostelium* | * *Coremiostelium polycephalum* |
| | *Dictyostelium polycephalum* | Dpol | B1Aa | | * *Coremiostelium polycephalum* |
| | *Dictyostelium polycephalum* | Dpol | 2A2j | | * *Coremiostelium polycephalum* |
| | + *Dictyostelium polycephalum* | Dpol | 1JNd1 | | * *Coremiostelium polycephalum* |
| Group 2B | + *Polysphondylium filamentosum* | Pfil | SH1 | Genus *Heterostelium* | * *Heterostelium filamentosum* |
| | + *Polysphondylium pallidum* | Ppal | 1U4Aa | | * *Heterostelium pallidum* |
| | *Polysphondylium pallidum* | Ppal | 2F2Ca | | * *Heterostelium pallidum* |
| | *Polysphondylium pallidum*[e] | Ppal | PN500 | | * *Heterostelium pallidum* |
| | + *Polysphondylium album* | Palb | 1X2Ca | | * *Heterostelium album* |
| | + *Polysphondylium tenuissimum* | Pten | E1M4 | | * *Heterostelium tenuissimum* |
| Group 2A | + *Acytostelium leptosomum* | Alep | FG12A | Genus *Acytostelium* | *Acytostelium leptosomum* |
| | + *Acytostelium subglobosum* | Asub | LB1 | | *Acytostelium subglobosum* |
| | *Acytostelium subglobosum* | Asub | 2CBg | | *Acytostelium subglobosum* |
| | + *Acytostelium amazonicum* | Aama | 2B1Ba | | *Acytostelium amazonicum* |
| polycarpum | + *Dictyostelium polycarpum* | Dpcp | S570 | Genus *Synstelium* | * *Synstelium polycarpum* |
| | *Dictyostelium polycarpum* | Dpcp | 1WC9Ap | | * *Synstelium polycarpum* |
| Group 1 | + *Dictyostelium fasciculatum* | Dfas | SH3 | Family Cavenderiaceae | * *Cavenderia fasciculata* |
| | *Dictyostelium fasciculatum* | Dfas | Smok | | * *Cavenderia fasciculata* |
| | + *Dictyostelium myxobasis* | Dmyx | 2D2Bb | | * *Cavenderia myxobasis* |
| | *Dictyostelium aureo-stipes* | Da-s | D2LKa1 | | * *Cavenderia aureostipes* |
| | + *Dictyostelium aureo-stipes* | Da-s | 1WC11Ab | | * *Cavenderia aureostipes* |
| | + *Dictyostelium granulophorum* | Dgra | CHII-4 | | * *Cavenderia granulophora* |
| | + *Dictyostelium microsporum* | Dmic | Hag143 | | * *Cavenderia microspora* |
| | + *Dictyostelium deminutivum* | Ddem | M19A | | * *Cavenderia deminutiva* |

**S1 Table. Species (strains/isolates) studied in this work.**

(a) Traditional species names used in this report. + : representative isolates of the 36 species shown in Fig 3, Fig 8, and S6 Table. (b) Four-letter abbreviations for use in figures and tables. They consist of the first letter of the genus and mostly the first three letters of the specific epithet. (c) Corresponding taxon names according to the new classification by Sheikh *et al*. [1]. Asterisks: species names that are different from the older names. (d) Possibly *D. clavatum,* according to the SSU rDNA sequence. (e) Possibly *P. album* (*H. album*)*,* according to the SSU rDNA sequence.

(See additional file “S2_Table.xlsx”)

**S2 Table. Location and climate classification of isolation sites.**

(a) The second set of coordinates is for the collection site of which the location could not be precisely specified. The location of isolation site was assumed to be within a rectangular area defined by the two sets of longitude and latitude that contains the main part of the place name shown in the source reference. The range of inaccuracy due to this method is not large, as shown in Fig 5. (b) Köppen-Geiger climate classifications [21]. (c) KI: K. Inouye, TU: T. Urai, IS: I. Shibano-Hayakawa, TM: T. Muranaka, c: laboratory course for undergraduate students organised by KI. (d) NBRP: National BioResource Project Cellular slime moulds, Japan (https://nenkin.nbrp.jp/), DSC: Dicty Stock Center (http://dictybase.org/StockCenter/StockCenter.html), ATCC: American Type Culture Collection (https://www.atcc.org/). (e) Designated as “Hagiwara 5” in [2].

| Species | Culture method |
|---|---|
| *D. septentrionalis* | Shaking |
| *D. leptosomum* | Plate |
| *D. implicatum* | Plate |
| *D. brefeldianum* | Plate |
| *D. capitatum* | Shaking |
| *D. rosarium* | Shaking |
| *D. sphaerocephalum* | Shaking |
| *D. mucoroides* | Shaking |
| *D. purpureum* | Shaking |
| *D. discoideum* | Shaking |
| *D. firmibasis* | Shaking |
| *D. giganteum* | Shaking |
| *D. laterosorum* | Plate |
| *P. violaceum* | Plate |
| *D. mocochasioides* | Shaking |
| *D. tenue* | Shaking |
| *D. minitum* | Plate |
| *D. rhizopodium* | Shaking |
| *D. vinaceo-fuscum* | Shaking |
| *D. lacteum* | Shaking |
| *D. caveatum* | Plate |
| *D. polycepualum* | Plate |
| *P. filamentosum* | Plate |
| *P. pallidum* | Plate |
| *P. album* | Plate |
| *P. tenuissimum* | Shaking |
| *A. leptosomum* | Plate |
| *A. subglobosum* | Shaking |
| *A. amazonicum* | Shaking |
| *D. polycarpum* | Plate |
| *D. fasciculatum* | Shaking |
| *D. myxobasis* | Plate |
| *D. aureo-stipes* | Plate |
| *D. microsporum* | Plate |
| *D. granulephorum* | Shaking |
| *D. deminutivum* | Shaking |

**S3 Table. Preculture method for each species.**

Shaking: cells were grown with bacteria in shaking culture. Plate: Cells were grown on LP agar plate with bacteria.

(See additional file “S4_Table.xlsx”)

**S4 Table. Monthly average temperatures at the collection sites of the 58 strains/isolates.**

The strains/isolates with three sets of 12 months are those of which the isolation site could not be precisely specified. They are median, minimum, and maximum temperatures for each month of the 0.5’ grid cells within the rectangular area defined by the two sets of longitudes and latitudes given in S1 Table. The strains/isolates are arranged in the order of phylogeny with group 4 at the top. Species names are shown in 4-letter abbreviation as listed in S1 Table.

(See additional file “S5_Table.xlsx”)

**S5 Table. Relative growth rate data of the 58 strains/isolates.**

From left, relative growth rates at 0 to 37 degrees celsius, mean, S.D., skewness, optimal growth temperature, and lower and upper temperature limits for 10%, 50%, and 90% maximal growth. The order of arrangement is the same as Table S2. Species names are shown in 4-letter abbreviation as listed in S1 Table.

(a) Entire tree (36 species)

| | mean | *P* | lower limit | *P* | upper limit | *P* |
|---|---|---|---|---|---|---|
| *lambda* | 0.5781 | 0.0057 | 0.5824 | 0.0029 | 0.4319 | 0.1659 |
| *K* | 0.2590 | 0.0074 | 0.2505 | 0.0109 | 0.2612 | 0.0186 |
| *I* | 0.3539 | 0.0010 | 0.3089 | 0.0010 | 0.2167 | 0.0010 |

(b) Branch I (14 species)

| | mean | *P* | lower limit | *P* | upper limit | *P* |
|---|---|---|---|---|---|---|
| *lambda* | 0.0001 | 1.0000 | 0.0001 | 1.0000 | 0.0001 | 1.0000 |
| *K* | 0.1124 | 0.6406 | 0.1224 | 0.5671 | 0.1151 | 0.6187 |
| *I* | -0.3567 | 1.0000 | -0.3611 | 1.0000 | -0.2164 | 1.0000 |

(c) Branch II (22 species)

| | mean | *P* | lower limit | *P* | upper limit | *P* |
|---|---|---|---|---|---|---|
| *lambda* | 0.7486 | 0.0010 | 0.7576 | 0.0001 | 0.5718 | 0.0797 |
| *K* | 0.7322 | 0.0005 | 0.6902 | 0.0012 | 0.5285 | 0.0102 |
| *I* | 0.6021 | 0.0010 | 0.6354 | 0.0010 | 0.3212 | 0.0010 |

**S6 Table. Phylogenetic signal metrics for the mean, lower limit, and upper limit growth temperatures using Pagel's *lambda*, Blomberg's *K* and Moran's *I*.**
In these indices, random changes of the traits over evolution give values close to one, whereas diversification under strong selection pressure would give a value close to zero. The values computed for the entire tree and separately for the two branches of the tree are shown. Branch I is made up of groups 1, 2A, 2B, plus one small clade, and Branch II, groups 3 and 4, plus two small clades. *P*: *P*-value for the null hypothesis of no phylogenetic signal.

### S1 Appendix

#### Calculation of the optimal, mean, and lower/upper limits of growth temperature

Some of the measures of thermal characteristics were defined and calculated from the growth rate vs. temperature (G-T) relationship using its 1st, 2nd, and 3rd order moments ($M_1$, $M_2$, $M_3$, respectively) for each strain/isolate.

$$M_n = \frac{\Sigma t^n \cdot g(t) \cdot \Delta t}{\Sigma g(t) \cdot \Delta t}$$

$$V = M_2 - M_1^2$$

$$S = M_3 - 3 \cdot M_1 \cdot M_2 - M_1^3$$

where $M_n$ is *n-th* moment, $V$ variance, $S$ skewness, $t$ temperature, $g(t)$ growth rate at $t$, and $\Delta t$ step size, which are 4, 5, 6, 6, 6, 4.5, 3, 3, and 3 from 0 to 37°C.

Optimal growth temperature (OGT) was estimated as the average of LL90 and UL90 which had been corrected for the asymmetry of the G-T relationship by its skewness.

$$OGT = \left(LT90 \cdot (1 - S) + UT90 \cdot (1 + S)\right) / 2 \quad (1)$$

Mean growth temperature was defined as the 1st order moment of the G-T relationship ($M_1$). For the lower and upper temperature limits for growth, temperatures giving 10% maximum growth rate were estimated using the moments of the G-T relationship (LL10 , UL10).

$$LT10^* = M_1 - 2 \cdot V^{1/2} + 3 \cdot S$$

$$UT10^* = M_1 + 2 \cdot V^{1/2} + 3 \cdot S$$

These values were used because LL10 and UL10, which are directly obtained from the G-T relationship, are sensitive to small differences in the growth rate value. The coefficients of the terms (2 and 3) were empirically determined.
